# Breakups and Hookups: a Markov model for karyotype evolution

**DOI:** 10.1101/2023.08.15.553394

**Authors:** Derek Setter

## Abstract

Chromosome rearrangements represent a prominent form of genetic variation that plays a key role in creating genetic isolation between emergent species. Despite their significance, the mechanisms and constraints governing chromosome evolution remain poorly understood. Relatively few species have karyotypes with very high chromosome counts, and the chromosome sizes of most species tend to be narrowly distributed around the mean length. Here, we develop and analyze a Markov model for the evolution of chromosome number and relative sizes through fission and fusion events, exploring several alternative models for the dynamics of each as well as the effect of enforcing limits on chromosome length. We compare the distribution of chromosome lengths predicted by the Markov model to karyotype data for a range of Eukaryote species to identify the best-fitting fission/fusion dynamics. We find broad support for a model which (i) favours the breaking of long chromosomes, (ii) favours the fusion of pairs of small chromosomes, and (iii) does not require size limitations to provide a good fit to the data. However, there are exceptions. On the one hand, species with micro chromosomes fit best to models with more uniform rates of fission and/or fusion. On the other hand, many species have chromosome sizes that are much more narrowly distributed than our models predict, suggesting the need to explore alternative dynamics and/or limitations to chromosome lengths.

## Introduction

Chromosome rearrangements are perhaps the most conspicuous type of genetic variation. Karyotypes are known to differ among closely related species and to play a role in reinforcement of both pre-zygotic and post-zygotic barriers to gene flow between emerging species, including mice (Hauffe and Searle, 1998; Bidau et al., 2001), nematodes (Yoshida et al., 2023), buttercups, (Baltisberger and Hörandl, 2016), and butterflies (Lukhtanov et al., 2005; Mackintosh et al., 2023a). While some clades, such as Lepidoptera, have highly conserved chromosomes dating back hundreds of millions of years (Wright et al., 2023), karyotype can also evolve very quickly, e.g. the muntjack deer has experience 26 chromosome fusions over the course of just a few million years. (Mudd et al., 2020). Although karyotype evolution can be rapid, there are clear limits on chromosome rearrangement. Most species have relatively few chromosomes and the distribution of chromosome sizes tends to be quite narrow (Li et al., 2011). However, despite the ubiquity and importance of karyotype differences among species, surprisingly little known about mechanisms and limits that underlie karyotype evolution, in part due to a lack of sufficient theoretical models.

Differences in chromosome number *k* are generated by two types of structural rearrangements: fission increases *k* by breaking a chromosome into two, while fusion decreases *k* by joining two separate chromosomes together end-to-end. This contrasts with translocation in which part of a chromosome is transferred to another, possibly reciprocally, such that two chromosomes swap segments without changing *k*. On the one hand, models of reciprocal translocation (Sankoff and Ferretti, 1996; De et al., 2001) have focused on understanding the distribution of chromosome lengths observed within species. On the other hand, models of fission and fusion (Yoshida and Kitano, 2021; Hipp, 2007) have focused on the evolution of chromosome number but disregard size.

Sankoff and Ferretti (1996) previously developed a simple Markov model for the evolution of chromosome size through reciprocal translocation. In their model, the genome is distributed among *k* chromosomes with different sizes, and at each step, two chromosomes are chosen to swap genetic material at break point distributed uniformly along their lengths. They considered two models: one in which chromosomes are chosen with equal probability and one in which they are chosen with probability proportional to the size. They compiled chromosome data for several eukaryotic species and tested the fit of their model to the observed distribution of chromosome sizes. Proportional break rates provided a better fit for many species, including Zea mays, Wheat, and Humans. In general, however, observed chromosome sizes have a much more narrow distribution than their model predicts. For this same data, De et al. (2001) show that accounting for the presence of centromeres by allowing translocation events to occur among chromosome *arms* improves the fit. However, in both studies, size constraints were needed in order to accurately predict chromosome sizes in these species. Indeed, Li et al. (2011) found that across many eukaryote species chromosome sizes tend to be centered around the average length and that there is a widely-conserved boundary with minimum and maximum sizes of approximately 0.4 and 1.9 times the average. However, even when including these boundaries in the model, they show that the reciprocal translocation generates a distribution of chromosome lengths that is still too broad.

The poor performance of the reciprocal translocation model is likely due, at least in part, to the assumption of a fixed chromosome count. To be applicable, it requires that translocation events occur at a much higher rate than fissions and fusions so that the distribution of chromosome sizes returns to equilibrium after one of these two events. However, this would entail frequent and major changes in synteny, which is generally not observed, even in Lepidopteran species with long-term stable chromosome number (Wright et al., 2023). As such, it is clear that in most species, the distribution of not only chromosome number but also chromosome size is primarily determined by fission and fusion events.

To this end, Yoshida and Kitano (2021) developed a Markov model for the evolution of chromosome number through fission and fusion events that disregards chromosome size such that the break rate is proportional to chromosome number *k* and the fusion rate is proportional to 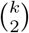. In addition, chromosomes are classified by their centromere location as *metacentric* (in the middle of the chromosome) or *acrocentric* (at the periphery). Only metacentric chromosomes can break (with equal probability), only pairs of acrocentric chromosomes may fuse (also with equal probability), and chromosomes can transition between these two states. However, chromosome size is not considered. They found that a narrow distribution of chromosome number is expected under this model and performed a phylogenetic analysis to show that evolutionary changes in the underlying rates of fusion and/or fission is required to explain karyotype varaition both between and within two different clades of fish. Hipp (2007) found a similar result in sedges using a Brownian motion model to explain variation in chromosome number. However, neither of these models make predictions about chromosome size and therefore cannot be tested directly against chromosome size data as was done for the analysis of the translocation model described above.

In this paper, we study a Markov model for the evolution of **both** chromosome number and size through fission and fusion events, proposing several different models for the dynamics of each process, here, disregarding centromeres. We evaluate the performance of our model by comparing predicted chromosome size distributions to three empirical data sets. (i) We use the autosome length data for a phylogenetically diverse set of eukaryotic species compiled by Li et al. (2011). This serves as a broad test for the general fit of our model and allows us to compare the performance to that of the translocation-only model. We refer to this as the *eukaryote* data. (ii) We also test our model against chromosome sizes for a subset of Lepidoptera species, combining the broad sampling of UK Lepidoptera species of Wright et al. (2023) obtained by the Darwin Tree of Life project with the Heliconious specific data from Cicconardi et al. (2021). These data serve several different and important purposes. Firstly, Lepidoptera are holocentric, meaning that microtubules bind to many kinetochores along the entire length rather than at a single centromere (Schrader, 1935), thus better satisfying the simplifying assumptions in our model. Secondly, while the mechanisms underlying karyotype rearrangement may vary widely across eukaryotes, they are likely to be shared among species of this single clade. Thirdly, although many species maintain an ancestral karyotype with *k* = 31 chromosomes, holocentrism is thought to enable more rapid karyotype evolution via fusions and fissions (Jankowska et al., 2015). Indeed, some lineages have experienced a large number of rearrangements in recent history (Wright et al., 2023). Furthermore, mounting evidence suggests that fusion events in Lepidoptera tend to involve small chromosomes, in part motivating several of the fusion models we propose. In particular, Cicconardi et al. (2021) found that all ten fusions that are ancestral to the Heliconius clade were between one of the smallest chromosomes and a larger chromosome. **Note** that the sex chromosomes (Z/W or X/Y) are *excluded* from the data in all analyses and that throughout the paper “chromosomes” refers only to the *autosome* set of each species.

Combining various models for the dynamics of fusion and fission (described in Methods, below) with the data sets above, we address several questions: (i) For the various scenarios we consider, what is the relationship between the (relative) rates of fusion and fission and the distribution of the number of chromosomes we observe in a randomly-chosen karyotype? (ii) How are chromosome sizes distributed under the different models and which of these provides the best explanation for the data we observed? (iii) Does the size distribution for a sample with *k* chromosomes depend on the underlying rates of fission and fusion? (iv) Which model best describes chromosome sizes in the data and are explicit limits on chromosome length required to make sense of the data?

## Methods

### Model and notation

We consider a continuous time Markov process for the evolution of chromosome size and number through fission and fusion, exploring various model combinations for the underlying dynamics of the process through simulations. For all models, we use *β* and *ϕ* to represent intrinsic break and fusion rates. At any given time, there are *k* = 1, 2, … chromosomes with (ordered) sizes *c*_1_ *≤ c*_2_ *≤* … *≤ c*_*k*_ such that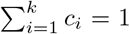, i.e. a genome of unit size. Throughout this paper, *ϕ* is set to one while the break rate is permitted to vary. In all models, for a fission event, a given chromosome breaks into two at a random site chosen uniformly along its length, while an end to end fusion of a pair of chromosomes creates a new chromosome that is the sum of their lengths. Below, we describe two distinct models of fission and three separate models of fusion, and we investigate the distribution of chromosome number and size under all six possible model combinations.

### Two models of breaking

#### Equal Break

In the first scenario, we treat fission as a meiosis-associated event in which any chromosome has an equal chance of breaking. Here, all chromosomes are assumed to break at the same rate *β* so that the total rate of fission events increases with chromosome number as *kβ*. We will refer to this as the Equal Break (EB) model.

#### Proportional Break

As an alternative scenario, we model fission as a mutation event that occurs at a genome-wide rate *β* and with an equal probability at every site. Here, the total break rate *β* is a constant value, i.e. independent of the number of chromosomes *k*, and the probability that any chromosome *i* breaks is proportional to its size *β ∗ c*_*i*_. We refer to this as the Proportional Break (PB) model.

### Three models of fusing

#### Equal Fuse

In the simplest model, any ordered pair of chromosomes fuses with an equal rate *ϕ*, such that the total fusion rate 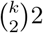*∗ ϕ* increases with the number of chromosomes. We refer to this as the Equal Fuse (EF) model.

#### Proportional Fuse

In the first alternative model, the rate that any (unordered) pair of chromosomes *i ≠ j* fuses is proportional to their inverse sizes: 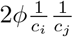. In other words, pairs of small chromosomes are much more likely to be involved in fusion events than pairs of large chromosomes. We refer to this as the Proportional Fuse (PF) model. Note that in this scenario, the total fusion rate is no longer a simple function of the number of chromosomes but instead depends on their relative sizes. The left panel of Fig. 1 illustrates the PF model for the *Bos taurus* karyotype.

**Figure 1:**
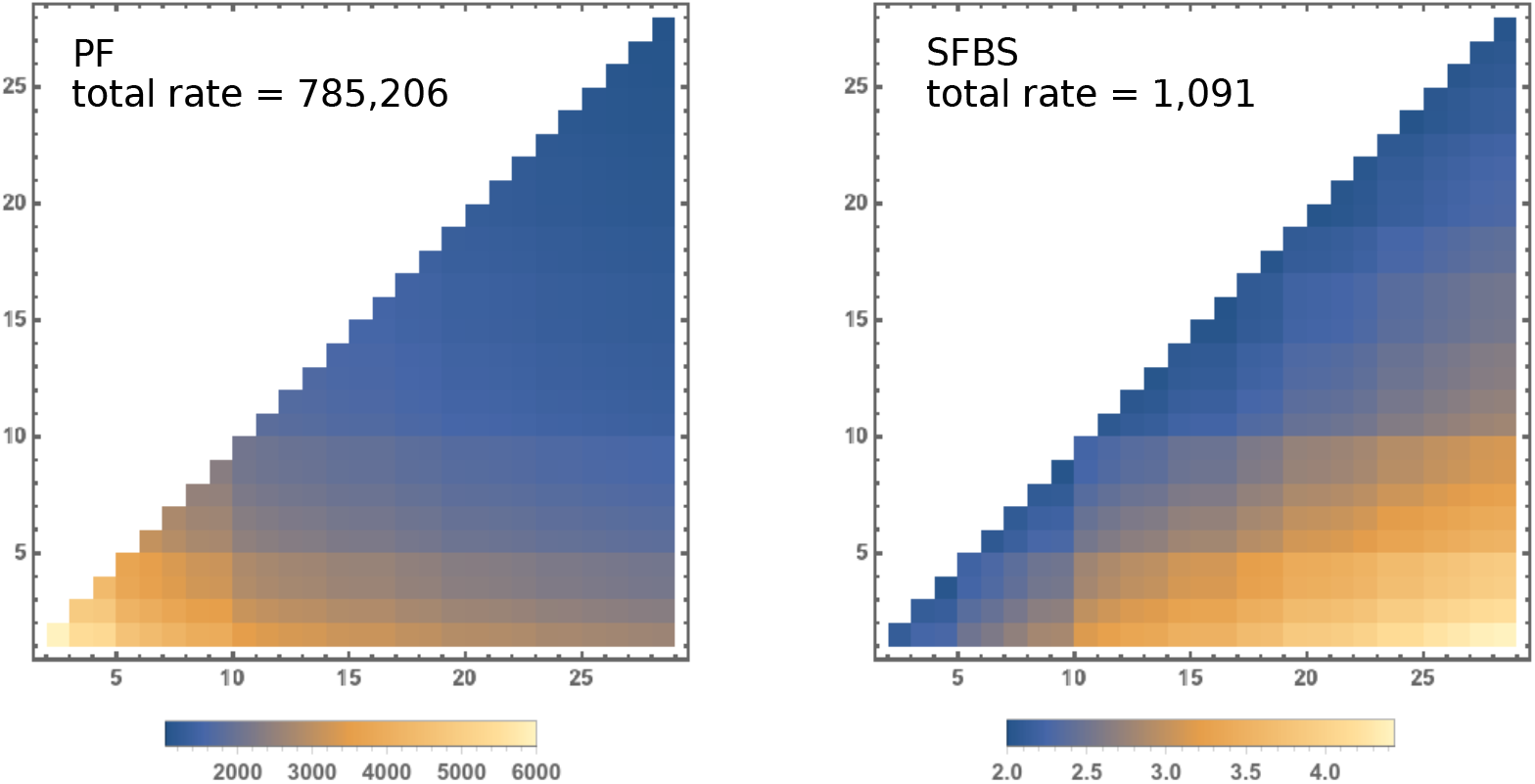
The rate at which each pair of chromosomes fuses under the PF model (left) and the SFBS model (right) in *Bos taurus*. The chromosomes are ordered from smallest to biggest on the x and y axes. The shading shows the relative rate of each events according to the respective scales below, and the total fusion rate is given for each of the models.

#### Short Fuse Big Stick

As a second alternative, we consider a scenario in which fusion events occur preferentially between one small and one large chromosome. Here, any (unordered) pair of chromosomes *i ≠ j* fuses with rate 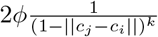, where the exponent *k* in the denominator ensures that the rates do not tend to 1 as *k* increases and the difference in chromosome sizes ||*c*_*j*_ *− c*_*i*_|| becomes negligible. We refer to this as the Short Fuse Big Stick (SFBS) model. As for the PF scenario, the total fusion rate at any point is no longer a simple function of *k*. The SFBS model is illustrated in the right panel of Fig. 1.

### Limitations on chromosome size

Consider the combination of the Equal Break and Equal Fuse (EB/EF) models. Here, small chromosomes have the opportunity to break into smaller and smaller pieces, and two large chromosomes have a high probability of fusing together. This contrasts with the clear limits on the minimum and maximum chromosome sizes observed in all data sets (Figure 2), particularly for Lepidoptera (middle and right panels).

**Figure 2:**
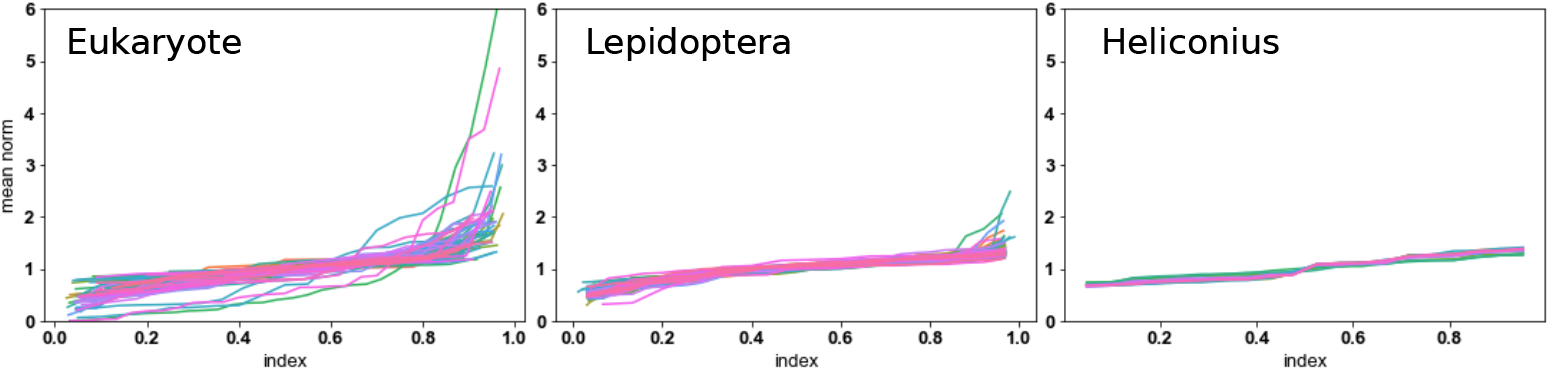
The distribution of relative chromosome sizes. Each line represents the data from a different species in the Eukaryote data (left), the Heliconius data (right), and the Lepidoptera DToL data (middle). The x-axis indexes the chromosomes from smallest to largest, placing them uniformly in the unit interval. For example, given *k* = 4 chromosomes, they are placed at positions 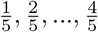. They y-axis shows the size of each chromosome relative to the mean.

One possiblity is to place a hard limit on chromosome size. Li et al. (2011) provide statistically motivated size boundaries derived from the eukaryote data common to our study (between 0.4 and 1.9 times the average). However, we found their size limitations to be too restrictive: 27 out of 73 eukaryote species and 6 out of 85 lepidoptera species have chromosome sizes outwith these limits.

Instead, we follow De et al. (2001) and apply a soft limit to chromosome size based on recombination rates and fitness. For successful meiosis, each chromosome must experience at least one recombination event. We assume that recombination occurs as a Poisson process at a per-base rate *r* such that that a chromosome with length *c*_*i*_ has a total recombination rate *c*_*i*_ *∗ r*. Without a loss of generality, we set *r* = 1 so that the probability a chromosome experiences at least one recombination event is 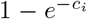. The fitness of an individual *u* = (*c*_1_, *c*_2_, …*c*_*k*_) with *k* chromosomes has fitness

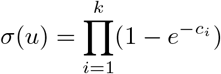

which is maximized when all chromosomes have the same length 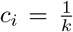. However, the maximum fitness that can be obtained depends on the number of chromosomes. In order to compare the fitness of individuals with differing numbers of chromosomes, we normalize by the maximum possible fitness:

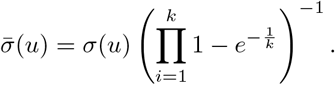

We use this relative fitness function to enforce a soft limit on chromosome size using the Metropolis algorithm. At a given step in the Markov process with *u* = (*c*_*i*_, …, *c*_*k*_) chromosomes, we simulate a step to new state *v* = (*c*_1_, …*c*_*j*_) with *j* = *k* + 1 or *j* = *k −* 1. Note that most but not all the *c*_*i*_ will be present in both *u* and *v*. If 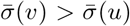, i.e. the fitness increases, we always accept the transition. When the relative fitness of the new state is lower, we accept the transition with probability 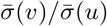, and otherwise, discard *v* and repeat this step until a successful transition occurs. We apply these soft limits to all six fission/fusion models and explore the effect this has on the distributions of chromosome number and size.

### Simulations

We investigate the distributions of chromosome number and size through Monte Carlo simulations of the Markov model under the various scenarios described above (see notebook S1 simulations.ipynb). The fusion rate is set to *ϕ* = 1 while the break rate *β* is allowed to vary. For any given *β*, the simulation begins with *k* = 1 chromosome. We run the simulations with an initial burn-in of 50,000 steps and sample the subsequent 100,000 consecutive steps. For a given state, we record the chromosome lengths and the exponentially distributed random holding time before transition to a new state occurs. The transition proceeds via a random fission or fusion event with a probability weighted by its relative rate. When the limits to chromosome size are enforced, transition events are smulated repeatedly until a successful fusion or fission event occurs. The holding time given that success occurs after *n* attempts is now the sum of *n* independent draws from the exponentially distributed holding time.

Chromosome number varies widely among the eukaryotes included in our data set (see Li et al. (2011)). Although the distribution in chromosome number will clearly depend on the underlying (relative) break rate *β*, there is currently no prior information about the rates associated with any given species or clade. We therefore use a two-step procedure to investigate the relationship between *β* and the expected number of chromosomes 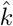 and to integrate over the uncertainty in *β* that is relevant for the data. In the first step, we test an initial range of *β* values to determine the relationship between *β* and 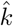. We then use linear interpolation to identify values *β*_2_, *β*_3_, … that approximately yield 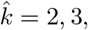, … chromosomes. We limit ourselves to a maximum of 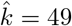 as this includes most of the species in our data set and has a reasonable computation time.

### Data

We explore three chromosome size data sets in our analysis:(i) A broad sampling of Eukaryote chromosome sizes compiled by Li et al. (2011) to test the fit of the reciprocal translocation model. We note that this data is older and may be imperfect, but it provides a direct comparison to their results. (ii) A broad sampling of the Lepidoptera species native to the UK obtained by Wright et al. (2023) for the Darwin Tree of Life project. Note that Lepidoptera are of course themselves eukaryotes and that this data set will be more complete and accurate than that of Li et al. (2011) (iii) A sample of 16 Heliconious species from within Lepidoptera obtained by Cicconardi et al. (2021).

## Results

In the main text, we will focus on four of the six models: Equal Break/Equal Fuse (EB/EF), Proportional Break/Proportional Fuse (PF/PF), Equal Break/Short Fuse Big Stick (EB/SFBS), and Proportional Break/Short Fuse Big Stick (PB/SFBS) as well as their complementary size-limited versions: EB/EF lim, PB/PF lim, EB/SFBS lim, and PB/SFBS lim. The EB/EF and PB/PF scenarios represent two ends of a spectrum in which fusion and fission rates depend on individual chromosome sizes: the former being the least intrinsically restrictive on chromosome sizes, while the latter, the most restrictive. The EB/SFBS and PB/SFBS models are chosen to highlight the effect of the alternative fusion that depends on the relative sizes of the two chromosomes. The complete model analysis is provided in the supplementary jupyter note-book *S2 model analysis*.*ipynb*, and the model fitting and goodness of fit tests can be found in supplementary notebook *S3 model fitting*.*ipynb*.

### Fragile genomes do not shatter

We first investigate the relationship between the break rate *β* and the mean number of chromosomes 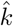 (Fig. 3). Under the EB/EF model, 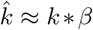, and this linear relationship is maintained under the EB/EF lim model with fitness-based limitations on karyotype evolution, though now with a slope closer to 1/2 (compare top left and top right panels of Fig. 3). In contrast, for the PB/PF and EB/SFBS, PB/SFBS models, Fig. 3 shows that a linear change in mean chromosome number for 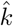 in the range of 2 to 49 requires non-linear magnitude-order changes in the break rate on the scale of (1e1 to 1e7) in the PB/PF model, (1e1 to 1e10) in the EB/SFBS model and (1e1,1e5) in PB/SFBS. While size limitations have little effect on this relationship for the PB/PF lim and PB/SFBS lim model, they shrink the scale over which *β* must vary in the EB/SFBS lim model: from approximately 1e1 for 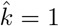 to 1e5 for 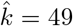. This suggests that proportional break and/or proportional fusion is sufficient to keep the chromosome size distribution near the optimum fitness.

**Figure 3:**
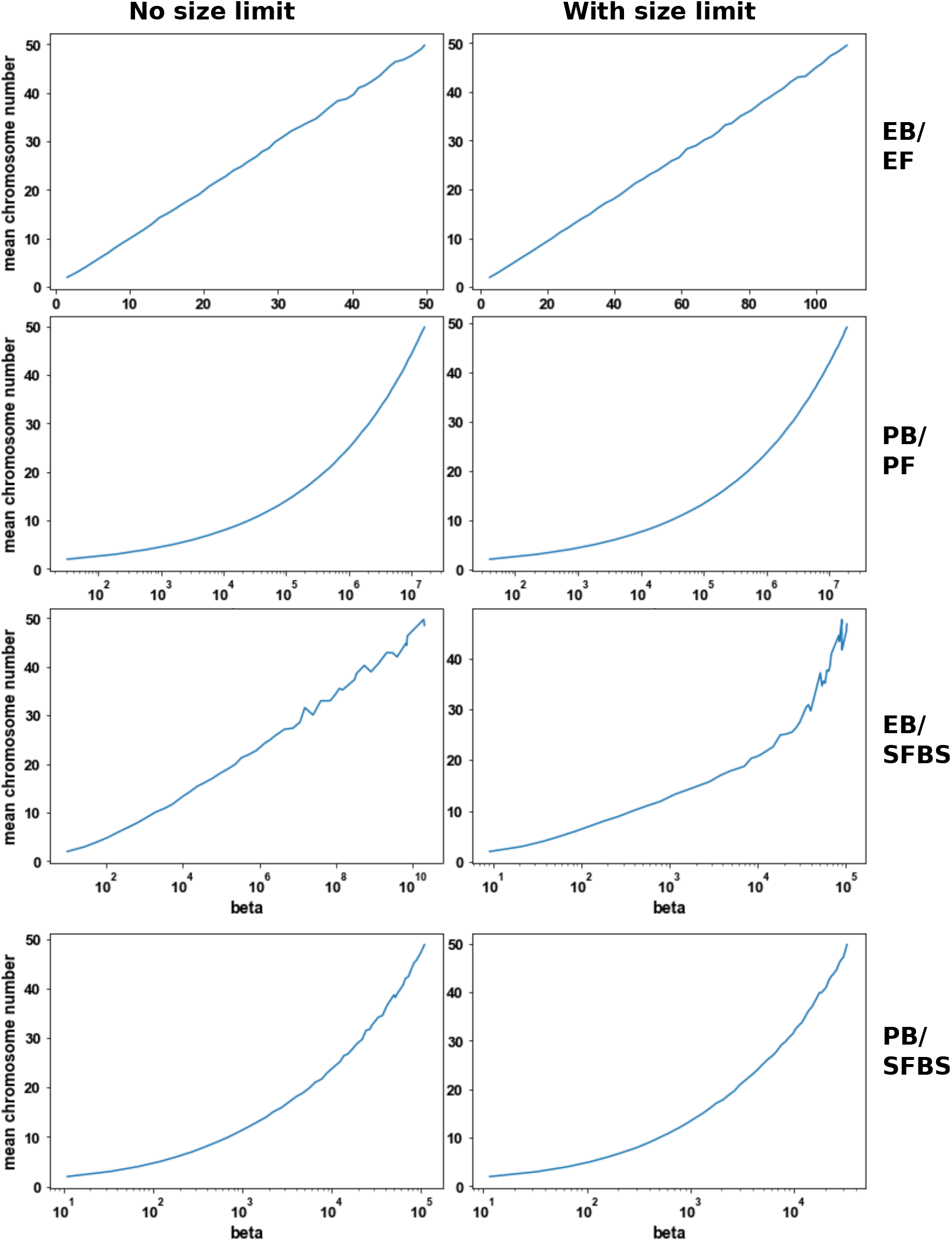
The relationship between the mean number of chromosomes observed 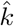 and the break rate *β* under three different models. From top to bottom, the rows correspond to four different models: EB/EF, PB/PF, EB/SFBS, and PB/SFBS. In the left column chromosome sizes are free to evolve; in the right, soft restrictions on chromosome sizes are enforced (see Methods). The y-axis in each panel is identical and shows the expected number of chromosomes for a given *β* on the x-axis. Note that the scale of the x-axis is unique to each panel: the EB/EF model is plotted on a linear scale in *β* while the PB/PF, EB/SFBS, and PB/SFBS models have a logarithmic scale on the x-axis.

Although there are many exceptions, a majority of species have karyotypes with fewer than 100 chromosomes. How do we interpret this in the context of these three scenarios? The EB/EF model suggests that most taxa have relatively similar break rates and that the generally tight range of chromosome numbers observed reflects a small upper bound to the relative rate that fission occurs per-chromosome. In contrast, under the PB/PF, EB/SFBS, and PB/SFBS models, the rate at which breaks occur (either per-site or per-chromosome, respectively) must vary widely among species. Although an upper bound on the break rate would likely exist, the general limit on chromosome number among species may in part reflect the diminishing returns for an increase in chromosome number with an increase in break rate.

### Stationary distribution of chromosome number

Above, we explored how the expected number of chromosomes 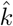 depends on the break rate *β* for the three focal scenarios. In this section, we investigate the full distribution of chromosome number under each scenario. As a first step, we derive the stationary distribution for chromosome number as a function of *β* and *ϕ* under the EB/EF scenario. Remember that in this scenario, the total rate of fission is *kβ* while the total rate of fusion is 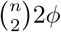.

Let *p*_*i*_(*t*) denote the probability that there are *i* chromosomes present at time *t*. The stationary distribution of this process can be obtained from the detailed balance equation. At stationarity, the change in probability *p*_*i*_ from *t* to *t* + *δt* is 0. That is, the rate of flux into and out of *p*_*i*_ is equal. Let *x*_*i*_ be stationary probability of being in state *i*. Let *λ*_*i*_ be rate of change from *i → i* + 1 and *µ*_*i*_ the rate of change from *i → i −* 1 chromosomes. At stationarity, the total rate of change is balanced, i.e.

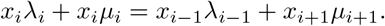

For the edge case,

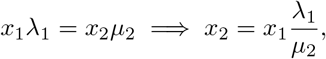

and generally,

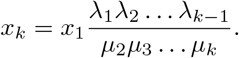

Allowing *λ*_*i*_ *→* (*i*)*β* and *µ*_*i*_ *→* 2*ϕ*(*i*)(*i −* 1)*/*2 as described above,

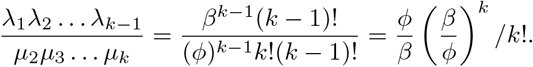

Furthermore, we know that 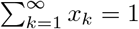, so that

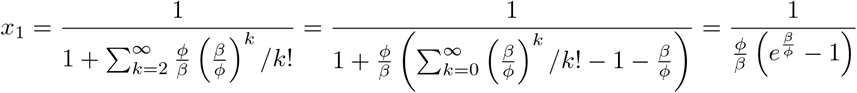

Note that this corresponds to the results of Yoshida and Kitano (2021) when centromeres are ignored. Let 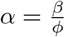. The mean and variance of the distribution are:

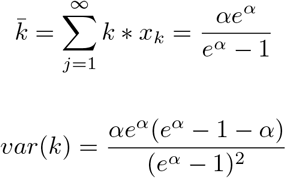

In Figure S1, we plot the mean and variance as a function of the relative break rate *β* with *ϕ →* 1 and observe that both the mean and variance converge to 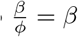 by the time *β* approaches around 8.0.

Figure 4 compares these analytic predictions to the realized distribution under each of the focal scenarios. First, focus on the scenarios with no limit on chromosome size (left column). Reassuringly, the simulated distribution under the EB/EF model closely matches the corresponding predictions derived above, with the distribution centering on the mean and a variance that increases with the mean. The PB/PF model has a distribution very similar to the EB/EF model except with a lower variance. In contrast, both the SFBS models have increased variance, and in combination with the EF model, results in a skew toward lower chromosome numbers and a heavy-tailed distribution.

**Figure 4:**
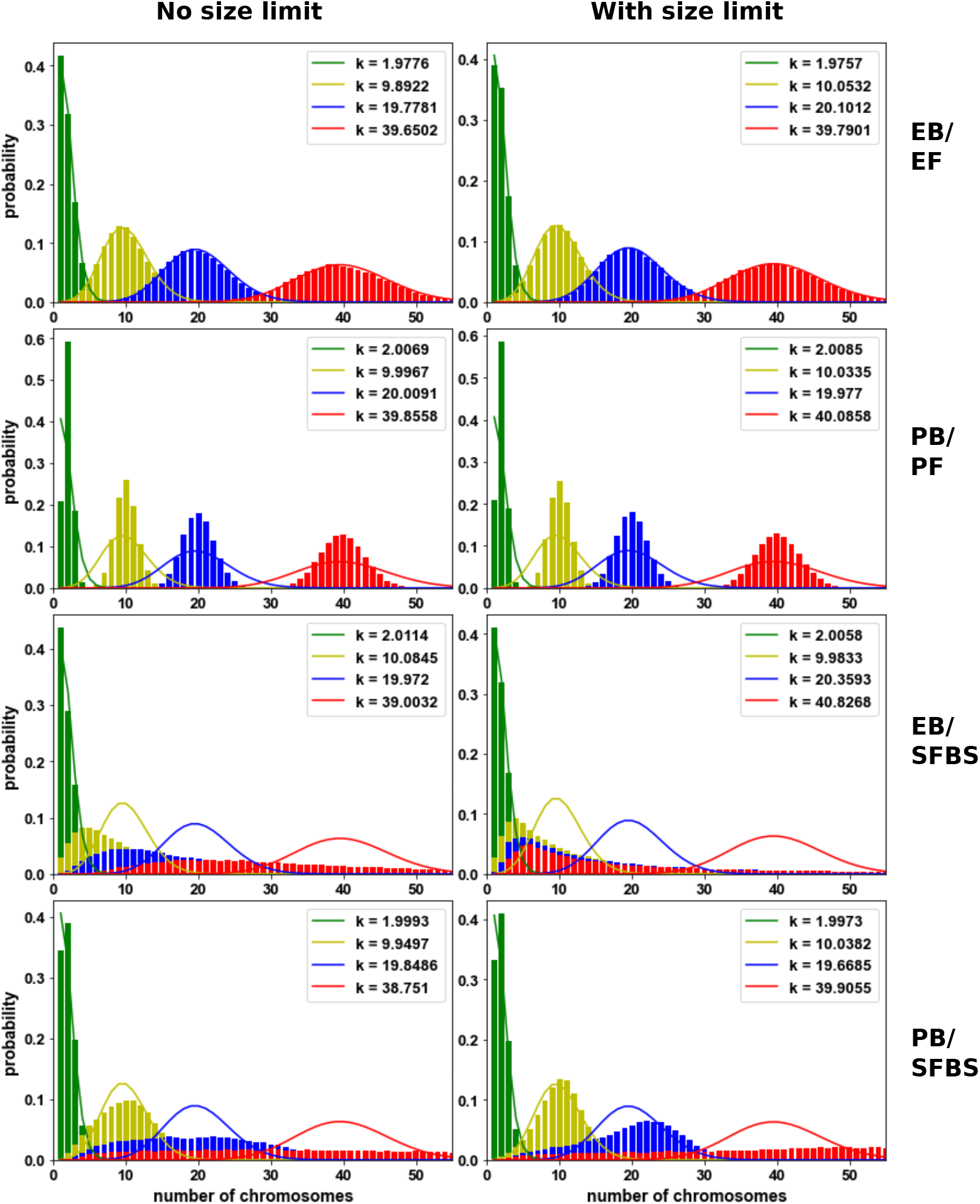
The stationary distribution for the number of chromosomes. The panels correspond to the same scenarios as in Fig. 3. For comparison across scenarios, the lines in each panel show the analytic predictions under the EB/EF model for 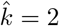 (green), 10 (yellow), 20 (blue), and 40 (red) (corresponding to break rates *β* = 1.59, 10, 20, and 40 respectively). The bar plot shows the scenario-specific distribution obtained for simulations with approximately the same corresponding 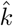 values. The legend shows the realized 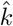 for each scenario. Note that the x-axis is the same in all panels but each has a unique y-axis.

The enforcement size limitations (compare left and right columns of Fig. 4) has little effect on the distribution of chromosome number in the EB/EF and PB/PF models. In contrast, the addition of size limits for the EB/SFBS model skews the distribution even more strongly toward smaller chromosome numbers. Surprisingly, the opposite is true for the PB/SFBS model: enforcing limitations skews the distribution toward larger chromosomes.

Chromosome number varies widely among eukaryotes and even between closely related species. The relatively narrow distribution in chromosome number observed for the PB/PF model (both with and without size limits) would suggest that lineages with very different chromosome numbers are likely to experience different break rates. This is true for all models we considered except for the SFBS models with no size limit (see *S2 model analysis*.*ipynb*). In these scenarios, a wide range of chromosome number may be observed among species with no need to invoke lineage-specific break rates.

### The chromosome size distribution

In this section, we describe the shape of the distribution of chromosome sizes under the various fission/fusion models and explore how this distribution depends on the underlying break rate *β*.

First consider the top left panel of Figure 5 corresponding to the EB/EF model. The white boxes in this plot show the distribution of chromosome sizes when *k* = 12 chromosomes are sampled from the simulation with a mean chromosome count of 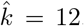. Teal shows the distribution for a sample of *k* = 12 from a simulation with 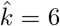 (i.e. we observe more chromosomes than expected in the simulation), while magenta shows the case when the simulated 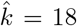 (i.e. we observe fewer chromosomes than expected). Figure 5 shows that, under the EB/EF model, there tend to be many small chromosomes and a few relatively large ones, and the distribution of chromosome sizes does not depend strongly on the underlying break rate. The addition of size limitations narrows this distribution by preventing tiny chromosomes from forming.

**Figure 5:**
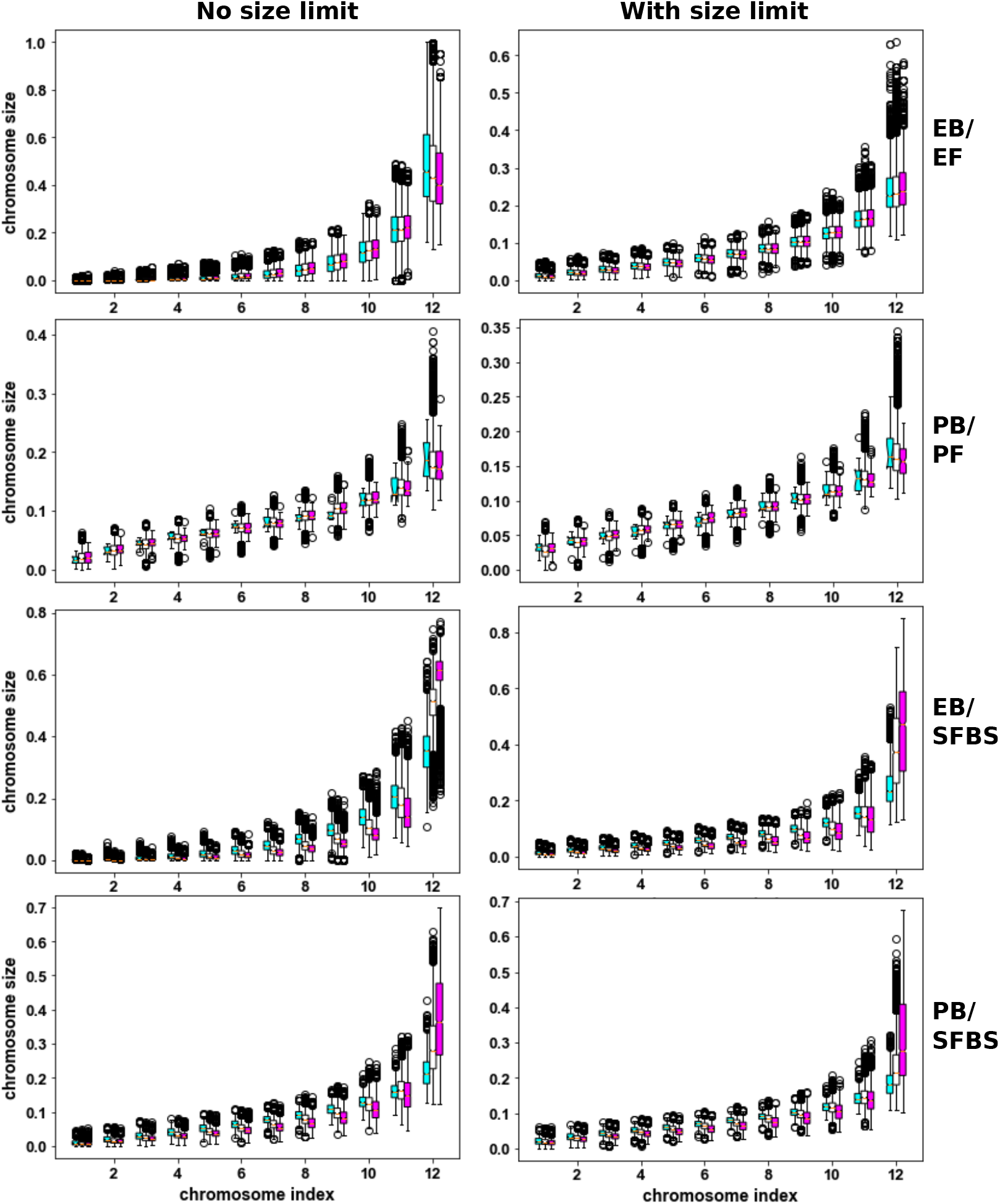
The chromosome size distribution and the dependence on the break rate. The panels correspond to the same scenarios as in Fig. 3. Each panel shows the distribution of (ordered) chromosome sizes when *k* = 12 chromosomes are sampled from simulations with different underlying break rates chosen such that 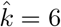 (teal), 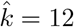 (white), or 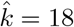 (magenta).

As expected, chromosomes sizes have a much narrower distribution under the PB/PF model (Fig. 5, second row). Here, the addition of size limitations does not have a significant effect, suggesting that the dynamics of the PB/PF controlling chromosome size help generate and maintain individuals with high fitness. As for the EB/EF model, there is not a significant affect of the underlying break rate. In contrast, the short fuse big stick model results in a broad distribution in chromosome sizes, similar to the EB/EF model. Indeed, the EB/SFBS model generates both the smallest and largest chromosome sizes among all models, while the distribution under PB/SFBS is more narrow than that of EB/EF. For both SFBS models, the underlying break rate has a strong affect. Given *k* chromosomes are observed, the higher the underlying break rate, the more likely it is to see very large and very small chromosomes.

### Proportional Break/Proportional fuse fits best

Here, we examine how well the six model combinations, both with and without size limitations, are able to explain the narrow distribution of chromosomes sizes observed across eukaryotes. To begin with, for a given model and focal chromosome number *k*, we extract from the simulations all instances where *k* chromosomes were observed, aggregating over the simulations with 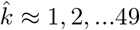, thus capturing as much variation in the chromosome size distribution as possible. The chromosomes are ordered from shortest to longest, and we calculate the mean size for each of the (ordered) chromosomes across all the sampled simulations. The fit of the model to a given species is measured by the sum of the square distance of the observed (ordered) chromosome sizes to the respective means. The best-fitting model is taken to be that with the smallest total square distance from the observed chromosome sizes.

We then perform a goodness-of-fit test for the best-fitting model of each species through bootstrapping, again using the aggregate data. For a given best-fitting model and species we obtain the total square distance from the mean for 1000 sub-samples from the corresponding simulation data and calculate a percentile score. This score represents the proportion of sub-samples with square distance less than that observed for the species such that a score of 1 indicates the worst-possible fit.

The best model for the vast majority of species examined in this study is the size-limited proportional break/proportional fuse model, PB/PF lim. Although this includes all Lepidopteran species, the model is a poor fit in this phylum. Only two species have goodness-of-fit scores within the 95% percentile: *Hedya salicella* (0.88) and *Notocelia uddmanniana* (0.92). Intriguingly, both of these species have a chromosome count that differs from the ancestral karyotype of *k* = 30. *H. salicella* has 24 chromosomes; *N. uddmanniana*, 27. Indeed, we can see that even the the PB/PF lim model predicts a chromosoem size distribution much broader than the narrow range of chromsome sizes observed in Lepidoptera species with the ancestral karyotype (Fig. 6).

**Figure 6:**
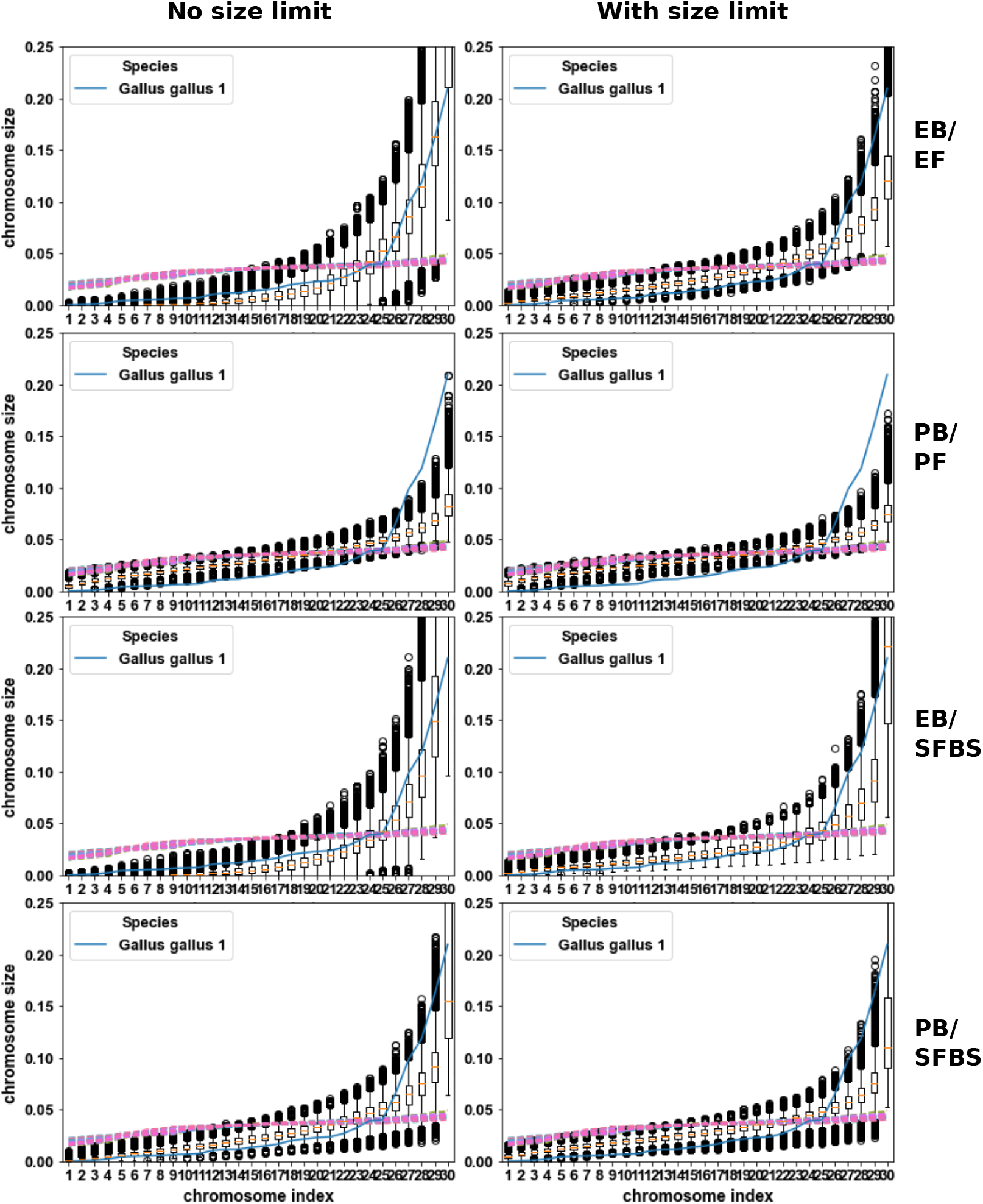
The size distribution when *k* = 30 chromosomes. The panels correspond to the same scenarios as in Fig. 3. The x-axis shows the chromosomes ordered from smallest (left) to biggest (right). The y-axis shows the distribution of the corresponding ordered sizes from the simulations, here, aggregating the samples where *k* = 30 over all *β* values simulated under the respective scenarios. The dashed lines correspond to the chromosome sizes observed for all Lepidoptera species with the ancestral karyotype of *k* = 30 chromosomes. Note that for better resolution, the y-axis is zoomed in to highlight the empirical observations and some simulation data may be cut off.

In contrast, the PB/PF lim model provides a good fit for many of the eukaryote species we examined. Figure 7 shows the range of percentile scores observed for the eukaryote species that fit the PB/PF lim model (blue). It compares it to the distribution of scores for the subset of 200 bootstrap samples that were both simulated under and best fit the PB/PF lim model (yellow). We see that many species have scores well within the distribution, however, there is an excess of percentile scores *>* 0.85. As for the Lepidoptera data, these represent species with chromosome sizes more narrowly distributed than expected under this model. Figure 8 shows several examples where this fission/fusion model fits the eukaryote data well and one example, *Glycine max*, where the model provides a relative poor fit (percentile score = 0.997).

**Figure 7:**
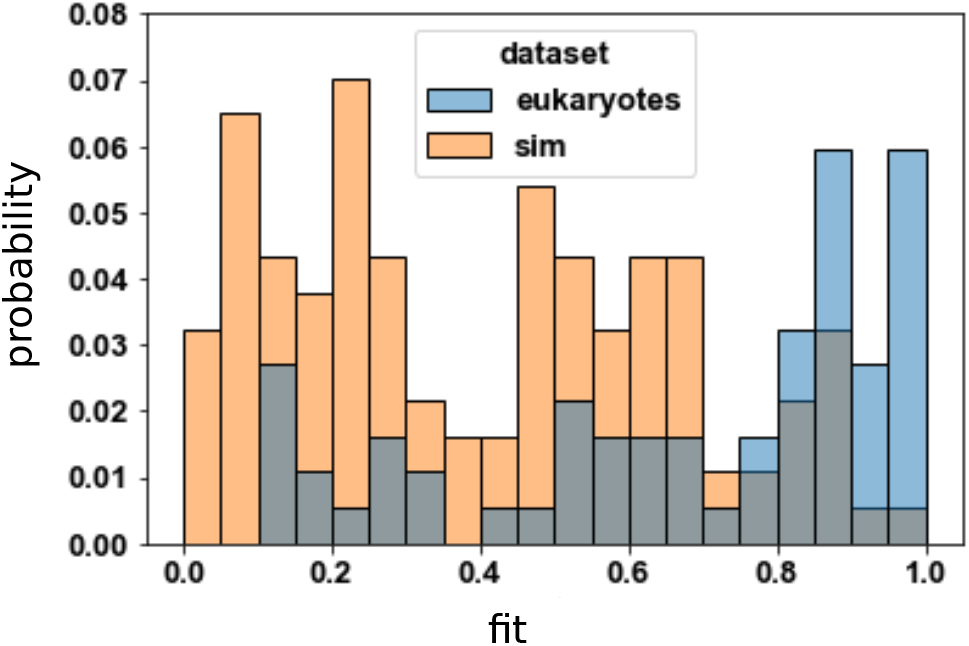
The distribution of goodness-of-fit scores for the PB/PF lim model. The y-axis shows the probability density for fitness scores in 20 bins evenly distributed between 0 and 1 (x-axis). The blue shows the percentile scores for eukaryote species (i.e. excluding Lepidoptera) for which PB/PF lim model provided the best fit. Yellow shows the percentile scores for simulation under the PB/PF lim model which were assigned the correct model.

**Figure 8:**
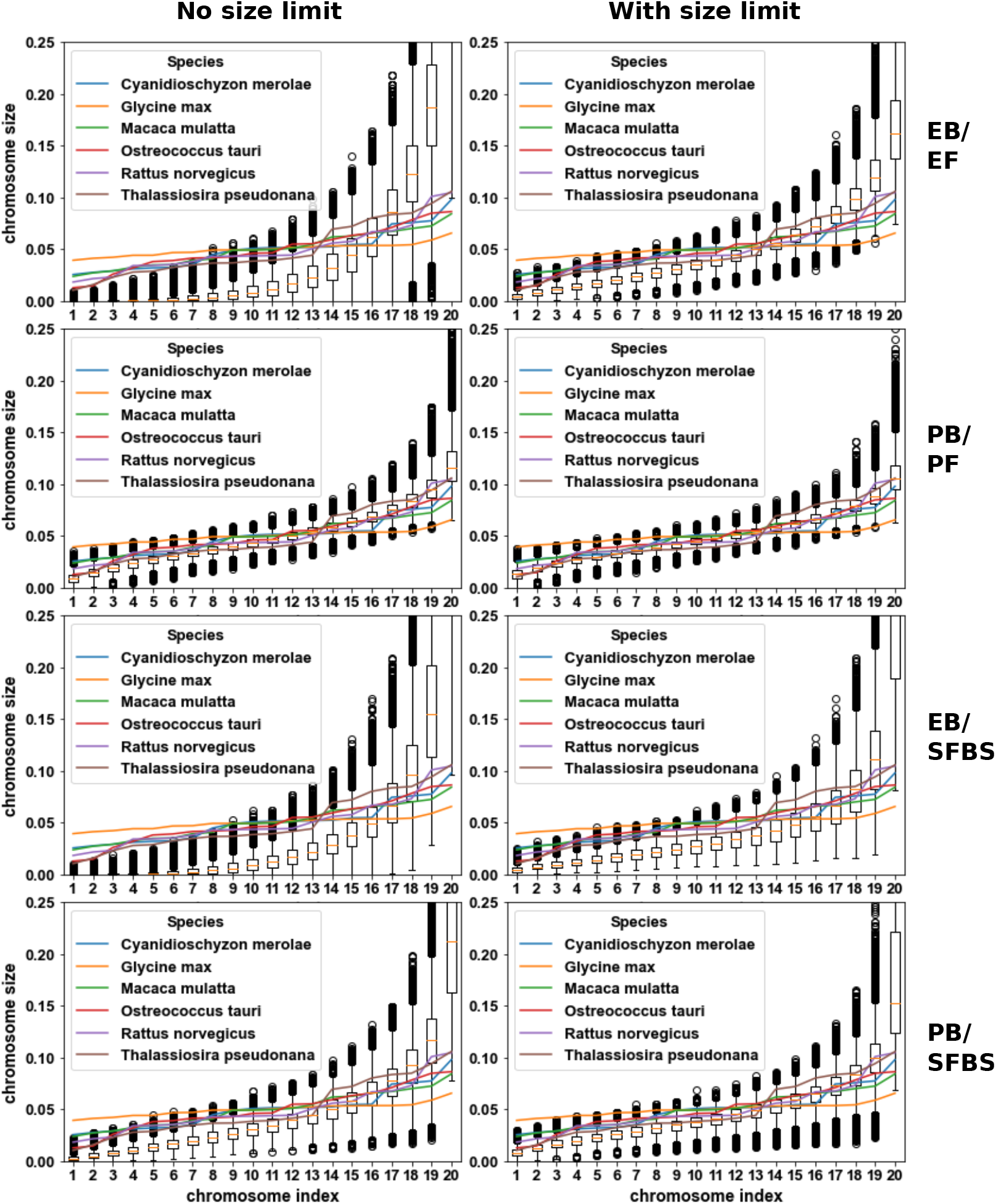
The size distribution when *k* = 19 chromosomes. The panels correspond to the same scenarios as in Fig. 3. The x-axis shows the chromosomes ordered from smallest (left) to biggest (right). The y-axis shows the distribution of the corresponding ordered sizes from the simulations, here, aggregating all examples where *k* = 19 over all *β* values simulated under the respective scenarios. Note that for better resolution, the y-axis is zoomed in to highlight the empirical observations and some simulation data may be cut off.

Table 1 lists the eight species that best fit one of the alternative models, noting any peculiarities about their genome organization and/or similarities among them. This suggests that non-standard recombination, ploidy, and sex chromosomes can affect the underlying dynamics of fusion and fission. Two of the outliers are both diatoms, suggesting differences in the dynamics of this clade compared to most eukaryotes. Perhaps reassuringly, both species in the dataset known to have microchromosomes fit a model that generates a broad distribution of chromosome sizes (the good fit of *G. gallus* to the EB/EF model can be seen in Fig. 6).

**Table 1:**
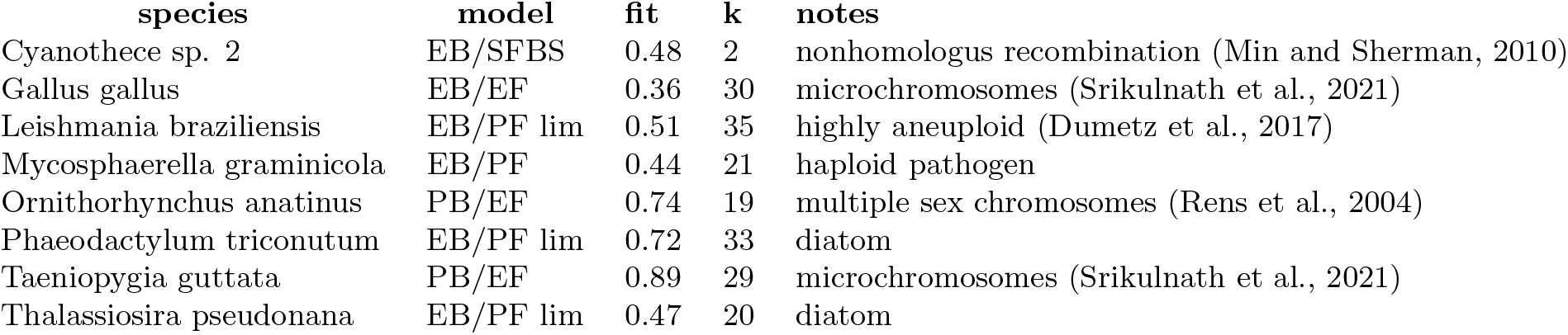
The best-fitting model, percentile score, chromosome count, and oddities about the eight species which did not fit the most-common PB/PF lim model.

One concern is whether we have power to discern between the different models given only the chromosome size data from a single species. To explore this, we calculated the confusion matrix for the assignment of each of the simulated data sets to the different models. For simulations with the most-common PB/PF lim model, Figure 9 shows there is relatively high power to correctly identify the model, but there is still a significant chance of wrongly classifying a species as following either PB/PF or EB/PF lim. This suggests that the few species we observed with the EB/PF lim model (*L. braziliensis, P. triconutum* and *T. pseudonana*) may in fact follow the dynamics of the PB/PF lim model. We found that both species with microchromosomes, *G. gallus* and *T. guttata*,follow an equal fuse (EF) model but differ in the break model. However, they both belong to the class Aves, and are not likely to differ in fission/fusion model. Indeed, Figure 9 shows it is possible that both species follow the EB/EF model but that *T. guttata* has been classified incorrectly.

**Figure 9:**
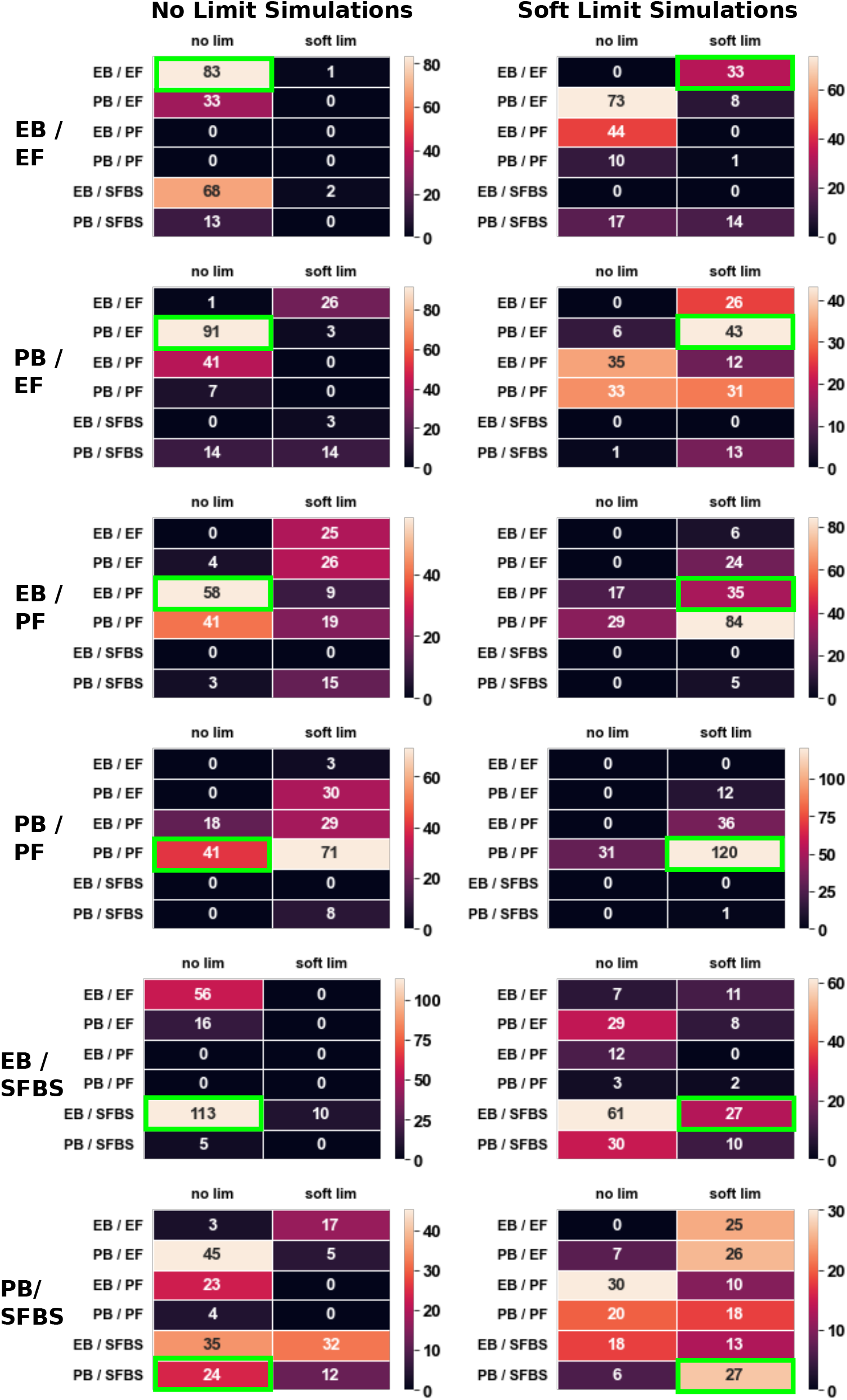
The confusion matrix for each of the simulated models. Each row corresponds to simulations under a combination of the fission/fusion models either with no size limit (left column) or with the soft limit on chromosome sizes (right column). A single plot corresponds to 200 simulations under the corresponding model and indicates the number of times each model was chosen as the best fit. The green box indicates the true underlying model.

## Discussion

Here we have explored several models for the fission and fusion of chromosomes to try to understand what dynamics underlie karyotype changes and how this affects the distribution of chromosome number and size. We find that a model with proportional break and proportional fuse (PB/PF) best fits a majority of the data. Surprisingly, this is true for all Lepidoptera species, despite the fact that they are holocentric (our model ignores centromeres) and that fusion events in these lineages tend to occur between small and large chromosomes (as in our Short Fuse Big Stick model). However, note that the fit is relatively poor, suggesting that these species follow some alternative dynamics of fusion and/or fission.

We find that there is only a slightly better fit of the PB/PF model when limits to chromosome length are enforced. This suggests that the general limit on chromosome number may be inherent to the dynamics of fission and fusion, as the break rate must increase orders of magnitude for a linear increase in chromosome number. Under the PB/PF model (and indeed all models we have considered except the size-unlimited Short Fuse Big Stick model), the distribution of chromosome number is relatively narrow, and large shifts in chromosome size within a clade are likely due to evolutionary changes in the underlying dynamics and/or rates of fusion and fission. While the PB/PF model provides a good fit to many eukaryote species, other taxa, particularly Lepidoptera, have a much more narrow range of chromosome sizes than this model predicts. This suggests the need to examine further alternative models for fusion and fission dynamics or more restrictive limits to chromosome length.

How do we interpret these results in a biological context? The EB model views fission as a per-chromosome event which implies that each chromosome has an equal chance of breaking during chromosome segregation in meiosis. In contrast, the PB model treats chromosome breakage as an event occurring at a uniform rate per-base in the genome, analogous to the process of random nucleotide mutations. As a consequence, larger chromosomes are more likely to break.

It is more difficult to place the chromosome fusion models in context. Our simplest model (EF) follows that of Yoshida and Kitano (2021), in which, any pair of chromosomes join end-to-end at an equal rate. The alternative fusion models seek to account for trends in the data. The proportional fusion (PF) model makes fusion between small chromosomes much more likely which generates a narrower distribution of sizes than the EF model. This could be the case if, e.g. the ends are more likely to be exposed in the 3-dimensional conformation of small chromosomes. In contrast, the SFBS model seeks to account for the observation that the divergence of *Heliconius* from the ancestral Lepidopteran karyotype resulted from the fusion of each of the 10 smallest chromosomes to one of the other 20 larger chromosomes.

### Not too big, not too small

The vast majority of species best fit the size-limited proportional break/ proportional fuse (PB/PF lim) model. However, in this scenario, restricting chromosome length mostly affects the variance and not the mean chromosome sizes. Indeed, among the unconstrained fission/fusion models, PB/PF is the most-likely model, and the inclusion of a size constraint provides only a marginal improvement in fit. This contrasts with the reciprocal translocation model that produces a reasonably good fit only when strict upper and lower bounds are applied (Li et al., 2011).

The size limits placed on our fission/fusion model differ from those of Li et al. (2011). They fit a gamma distribution to the eukaryote chromosome size data used in this study, and based on a 95 percent confidence interval, determined that most species have chromosome sizes between 0.4035 and 1.8626 times the mean. Using their limits, 26 of the 73 eukaryote species (common to our study and theirs) and 7 of the 84 Lepidoptera species (exclusive to our study) have chromosomes outside this range. Instead, we follow De et al. (2001) in applying a soft limit based on the probability that all chromosomes undergo a successful recombination event during meiosis. This represents a limit on chromosome size based on the change in fitness directly *caused* by a change in karyotype.

However, it is quite likely that evolutionary restrictions on chromosome size occur at the population level as a *consequence* of changes in karyotype. The models presented in this paper represent instantaneous changes in karyotype at the population level – equivalent to a substitution model in population genetics. In reality, a new structural rearrangement appears in a single individual and must spread through the population to reach fixation, and the relative contribution of natural selection and genetic drift to this process remains a matter of debate. These rearrangements will vastly and instantaneously change the physical association among large sets of loci in the genome that carry beneficial variation, and the fusion of chromosomes each carrying an adaptive allele will promote fixation of the rearrangement (Guerrero and Kirkpatrick, 2014). In addition, the landscape of recombination is altered (Bidau et al., 2001; Yoshida et al., 2023; Mackintosh et al., 2023a), e.g., assuming one cross-over per chromosome per meiosis, a shorter chromosome will have have a higher per-base recombination rate (Näsvall et al., 2023). This facilitates the dissociation and purging of deleterious variation and could promote fixation, particularly for fission of larger chromosomes. In contrast, and despite the pervasiveness of deleterious variation, in small and inbred populations, genetic drift may still be the predominant force (Wright, 1941; Lande, 1979). Indeed, Mackintosh et al. (2023b) find that, of the 12 chromosome fusions present in three closely-related species of *Brenthis* butterflies, only one shows evidence of historic adaptive evolution. Under genetic drift, the *physical* constraints outlined above will largely determine the distribution of chromosome sizes.

### One genome size does not fit all

One limitation of this study is that chromosome size is always expressed relative to the genome size, i.e. 0 *≤ c*_*i*_ *≤* 1. In reality, genome sizes vary greatly among different species and as a consequence, so do the physical lengths of chromosomes. It stands to reason that there is a lower limit on the physical length of a chromosome, e.g. Niwa et al. (1989) argue that for stable meiotic segregation in yeast, a chromosome has minimum size of 120 to 500kb including a functional centromere and telomere. In addition, physical constraints during meiosis may impose a strict upper limit on chromosome sizes. For very long chromosomes, the sister chromatids cannot fully separate before the daughter cells cleave apart, preventing successful meiosis. Schubert and Oud (1997) postulate that the longest chromosome arm cannot exceed half the average length of the spindle axis at telophase.

Although there are general trends in the distribution of (relative) chromosome sizes, the true physical length of a chromosome may not be the main factor that determines the rearrangement dynamics but rather its genomic content. Firstly, kinetochore length (and thereby strength of attachment of the centromere to the mitotic spindle) does not scale linearly with chromosome size (Plačková et al., 2022). Indeed, variation in centromere strength may influence which karyotype changes fix in a population through meiotic drive (Chmátal et al., 2014). Secondly, Peng et al. (2006) note that there is break-point re-use across clades that clusters in particular regulatory regions, purportedly in locations where breakage has minimal effect on gene expression. Thirdly, chromosome rearrangements frequently occur in regions with repetitive sequence, particularly transposable elements, that cause genome size expansion and promote ectopic recombination (reviewed in Li et al. (2017)). It is remarkable that the simple PB/PF model provides a good fit for many of the eukaryote species despite ignoring all of the biological details that we know to be important for karyotype evolution.

### Improvements and Prospects

We have already identified two limitations and potential improvements to the model, i.e. incorporating the true physical size of the chromosomes and identifying more biologically realistic fission/fusion models and chromosome size constraints. Here, we note several other aspects of the model that could be improved.

Anderson et al. (2020) demonstrate that pervasive sexual antagonism promotes the fusion of sex chromosomes and autosomes, particularly when diploid autosome numbers are low (*k <* 16). As such, the exclusion of sex chromosome dynamics in our model and data analysis is a glaring limitation. While the addition of sex chromosome fusion and fission dynamics are straightforward to incorporate into the model and could provide an avenue for future study, this process is more complicated than autosome-autosome fusions. The fusion of an autosome and sex chromosome generates a so-called *neo* sex chromosome and generally results in rapid evolution of gene content/expression (Zhou and Bachtrog, 2012; Bergero et al., 2015) and structural changes such as a reduction in chromosome size through degeneracy (Bachtrog, 2006). It will thus be important to identify a model that is both biologically realistic and analytically tractable.

The theoretical models thus far focus on either translocation or fission/fusion. However, it is clear that all three processes underlie the dynamics that shape karyotypes. Although such a complicated model may be analytically intractable even under the simplest of scenarios, it is straightforward to incoporate the translocation models of Sankoff and Ferretti (1996) and De et al. (2001) into our simulation framework and to include centromere effects as in Yoshida and Kitano (2021). It would be interesting to see if complex interactions arise from the interplay of the different types of chromosome rearrangement and whether this provides a more accurate explanation for the distribution of karyotypes observed across eukaryotes.

Finally, we have used the karyotypes of individual species to investigate the explanatory power of our fusion and fission models, but a phylogenetic approach may be more appropriate. For example, while the PB/PF model best fits the Lepidoptera species in this study, the predictions for chromosome size themselves are inaccurate. However, an alternative expectation maximization algorithm would likely support a SFBS model, particularly if applied to the rearrangement history ancestral to the *Heliconius* clade (Cicconardi et al., 2021). Indeed, a phylogenetic application of our model could prove particularly useful. In combination with the dependence of chromosome lengths on the underlying rates of fusion and fission, a phylogenetic approach would permit the co-inference of both the model and the underlying rates among among related lineages. Furthermore, current methods such as DESCHRAMBLER (Kim et al., 2017) and AGORA (Muffato et al., 2023) use maximum-parsimony to identify potential rearrangement histories among species but generally cannot determine the temporal sequence of events, and the most-parsimonious explanation may be neither unique nor correct. Our model could form the basis of a complementary likelihood-based approach for inferring historical karyotype changes.

## Acknowledgements

We would like to thank Alex Mackintosh and Konrad Lohse for helpful and insightful discussions as well as feedback on an early version of this manuscript. Support for the author was provided by an ERC starting grant (ModelGenomLand, 757648).

## Supportion Information

All supporting information is available at: https://github.com/dreq/Markov-Chromosomes.

## SI Figures

**Fig. S1:**
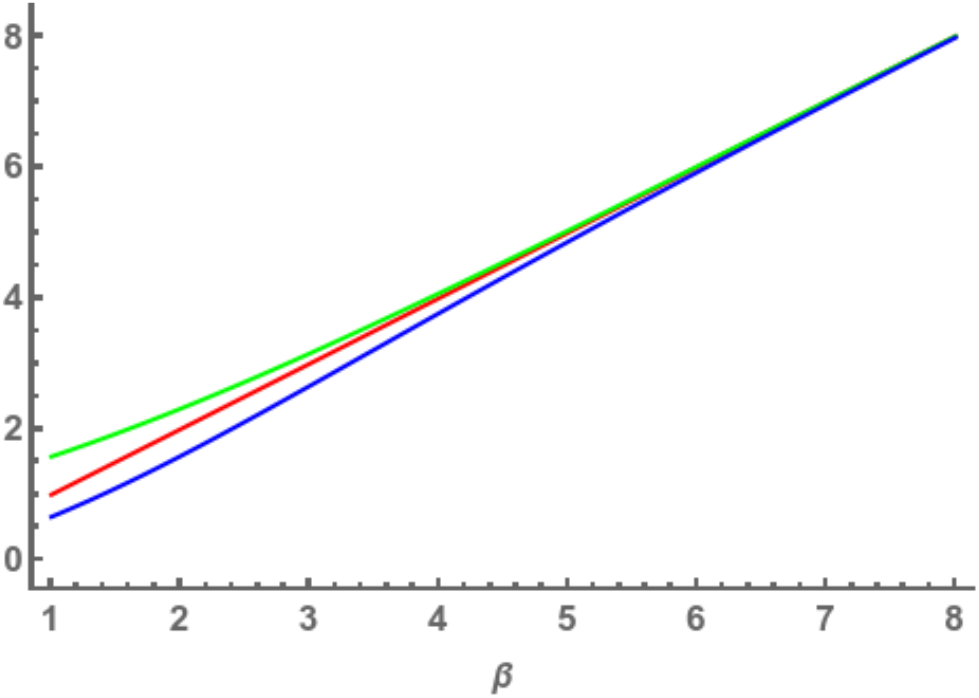
The mean (green) and variance (blue) for the chromosome number distribution under the EB/EF model. The function *y* = *β* is plotted in red to show the convergence of 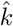 and *var*(*k*) to the break rate *β*.

